# baRulho: an R package to quantify degradation in animal acoustic signals

**DOI:** 10.1101/2023.11.22.568305

**Authors:** Marcelo Araya-Salas, Erin E. Grabarczyk, Marcos Quiroz-Oliva, Adrián García-Rodríguez, Alejandro Rico-Guevara

## Abstract

1. Animal acoustic signals are shaped by selection to convey information based on their tempo, intensity, and frequency. However, sound degrades as it propagates over space and across physical obstacles (e.g., vegetation or infrastructure), which affects communication potential. Therefore, transmission experiments are designed to quantify change in signal structure in a given habitat by broadcasting and re-recording animal sounds at increasing distances.
2. We introduce ‘baRulho’, an R package designed to simplify the implementation of sound transmission experiments. We highlight the package features with a case study testing the effects of habitat and acoustic structure on signal transmission. Synthesized sounds that varied in frequency, duration, and frequency and amplitude modulation were broadcast and re-recorded at five increasing distances in open and closed understory at the Bosque de Tlalpan, Mexico City. With this data, we showcase baRulho’s functions to prepare master sound files, annotate re-recorded test sounds, as well as to calculate and visualize measures that quantify degradation of acoustic signals in the time and frequency domain.
3. Degradation measures in baRulho adequately quantified acoustic degradation, following predicted patterns of sound transmission in natural environments. Re-recorded signals degraded less in open habitats compared to closed habitats, with higher-frequency sounds exhibiting more degradation. Furthermore, frequency modulated sounds degraded to a greater extent than pure tones. The increased attenuation and reverberation observed in higher frequency sounds and closed habitats suggest that factors such as absorption and scattering by vegetation play significant roles in transmission patterns.
4. The R package ‘baRulho’ provides an open-source, user-friendly suite of tools designed to facilitate analysis of animal sound degradation. Notably, it offers similar results to other sound analysis software but with significantly reduced processing time. Moreover, the package minimizes the potential for user error through automated test file annotation and verification procedures. We hope that baRulho can help enhance accessibility to transmission experiments within the research community, ultimately contributing to a deeper understanding of the ecological drivers of animal communication systems.

## 1. Introduction

Acoustic signals serve as a key means through which many animals convey information and navigate their social and ecological landscapes. Animal sounds are finely tuned by natural selection to maximize their effectiveness within specific ecological contexts (Bradbury & Vehrencamp 2011). This selective process, which acts on acoustic signals, is closely linked to the propagation of sound through natural environments and the challenges it entails. Transmission in natural settings can substantially impact signal integrity, potentially affecting their likelihood of detection and the successful transfer of information (Morton 1975; Marten & Marler 1977). As such, detailed insights into the complex interplay between animal acoustic signals and the environment, are critical to further our understanding of the mechanistic basis in the evolution of acoustic communication (Endler 1992; Cardoso & Price 2010; Tobias et al. 2010).

Sound transmission experiments are a key tool to evaluate the interaction of signals with a callers’ natural environment. Transmission studies seek to test hypotheses related to the degradation of sounds over space in combination with environmental factors that may influence selection on signal form and function (Graham et al. 2017; Benedict et al. 2021). Typically, this is achieved by broadcasting and re-recording animal sounds or synthesized sounds at increasing distances. Next, changes to the structural components of sounds are quantified by measuring modification to the power distribution in time and frequency domains and on combined time-frequency representations of sound (reviewed by Hardt & Benedict 2020). Such experiments have provided valuable insight into factors that affect signal transmission in natural environments. For instance, both habitat and acoustic structure can affect transmission in a significant manner (Kime et al. 2000; Apol et al. 2017; Wheeldon et al. 2022). In addition, other factors such as anthropogenic noise masking (Leader et al. 2005; LaZerte et al. 2015; Grabarczyk & Gill 2020), distance of the signaler from the ground, which leads to additional attenuation (Balsby et al. 2003; Darden et al. 2008; Arasco et al. 2022), as well as ambient temperature, humidity, and wind speed (Bradbury & Vehrencamp 2011), which also influence transmission patterns.

Conducting sound transmission experiments, however, can be challenging. Several steps are involved, such as careful formatting and manipulation of audio files, broadcasting and re-recording study signals in natural settings, annotation of re-recorded files, and quantification of degradation measures. These difficulties have likely contributed to the limited implementation of such experiments in animal communication research. Here, we introduce the R package ‘baRulho’ (Portuguese for ‘noise’), which is intended to facilitate animal sound transmission experiments and their subsequent analysis. The package offers tools to help researchers at each step of the process, from generating synthesized sounds and creating playback sound files, to streamlining annotation and measurement of sound degradation in re-recorded signals. We highlight package features with a case study testing the effects of habitat and signal structure on transmission properties using synthesized sounds. In addition, as proof of concept, we compared baRulho’s output to that of Sigpro (Pedersen 1998), the most commonly used software for quantifying animal sound degradation.

## 2. Implement transmission experiments with baRulho

Sound transmission experiments often follow a common sequence of stages (Table 1). The baRulho package offers a variety of functions to assist users during most stages. We showcase the package in an analysis workflow with our case study testing the effects of habitat and acoustic signal structure on sound transmission.

**Table 1.** Common steps in sound transmission experiments and associated functions in the baRulho package. For functions that compute degradation measures, which are listed under the ‘Quantify degradation’ stage, the “Description” field includes an explanation of the computed measure as well as selected citations to studies that employ the measure. Model sounds: sound in which degradation will be measured, usually found in the original field recordings or synthesized sound files. Reference sounds: sounds to use as a template to compare against for measuring degradation (typically re-recorded at 1 m). Test sounds: sounds re-recorded during playback experiments.

| Experimental stage | Function | Description |
| --- | --- | --- |
| Obtain signals / Synthesize sounds | `synth_sounds()` | Creates synthesized sounds that can be used for playback experiments. Can add variation to signal structure in five features: frequency, duration, harmonic structure, frequency modulation, and amplitude modulation. |
| Create master sound file for playback | `master_sound_file()` | Creates a sound file for playback experiments by clipping model sounds from one or more sound files and concatenating them in a single sound file, adding acoustic markers at the start and end to simplify time syncing of test sounds. |
| Time sync re-recorded sounds | `find_markers()` | Finds the time location of acoustic markers in the test sound files in order to time sync test sounds. |
|  | `align_test_files()` | Uses the time location of acoustic markers (determined by `find_markers()`) to determine the location of test sounds. |
|  | `auto_realign()` | Automatically adjusts the start and end of individual test sounds using spectrogram cross-correlation. |
|  | `manual_realign()` | Interactive graphic device to manually adjust the start and end of test sounds. |
|  | `plot_aligned_sounds()` | Creates image files with spectrograms to visually inspect alignment precision on test sound files (as single panels in Fig. 1). |
| Quantify degradation | `blur_ratio()` | Measures the mismatch between amplitude envelopes (expressed as probability mass functions) of reference and test sounds. By converting envelopes to probability mass functions the effect of power attenuation is removed, focusing the analysis on the modification of envelope shape. Higher values indicate more degradation. See also `plot_blur_ratio()` for visual inspection of blurring (Dabelsteen et al. 1993; Brown & Handford 2003, Fig. 3). |
|  | `envelope_correlation()` | Measures the similarity between reference and test sounds as the Pearson correlation between the two amplitude envelopes. Lower values indicate more degradation. |
|  | `excess_attenuation()` | Measures additional amplitude loss between reference and test sounds that is |
|  |  | greater than expected due to spherical spreading. Represents power loss due to additional factors like vegetation or atmospheric conditions. Higher values indicate more degradation (Holland et al. 1998; Darden et al. 2008). |
|  | *signal_to_noise_ratio()* | Measured as the ratio of sound power level of test sounds to that of ambient noise. Sound power measured as the root-mean-square of the amplitude envelope. Lower values indicate more degradation (Dabelsteen et al. 1993; Blumenrath & Dabelsteen 2004). |
|  | *spcc()* | Spectrographic cross-correlation between reference and test sounds as a measure of distortion in a time-frequency representation of sound. Computed as the maximum Pearson correlation from a sliding window procedure on Fourier or auditory scale spectrograms. |
|  | *spectrum_blur_ratio()* | Measures the mismatch between power spectra (expressed as probability mass functions) of reference and test sounds. Equivalent to *blur_ratio()* but in the frequency domain. Higher values indicate more degradation. See also *plot_blur_ratio(..., type = "spectrum")* for visual inspection of blurring. |
|  | *spectrum_correlation()* | Measures the similarity between reference and test sounds as the Pearson correlation between two power spectra. Equivalent to *spectrum_correlation()* but in the frequency domain. Lower values indicate more degradation. |
|  | *tail_to_signal_ratio()* | Measures the ratio of power in the tail of reverberations (found immediately after the sound) to that of the test sounds. Sound power is measured as the root-mean-square of the amplitude envelope. Higher values indicate more degradation (more reverberation) (Dabelsteen et al. 1993; Nemeth et al. 2006). |
|  | *plot_degradation()* | Creates multipanel plots (as image files) with spectrograms, amplitude envelopes, and power spectra of reference and test sounds, arranged by distance and transect to simplify visual assessment of degradation (as in Fig. 2). |

### 2.1. Case study: effects of habitat and signal structure on transmission

We conducted a sound transmission experiment to evaluate the effects of habitat and signal structure on degradation of synthesized sounds. Experiments were conducted on March 3, 2020 at Bosque de Tlalpan, Mexico City, an urban park composed of a mixture of xeric shrubland and oak forest. Reference sounds were recorded in open habitat (described below) at a mean sound pressure level of 82 dB, fast averaging, measured at 1 m from the playback speaker (Bose color soundlink). Test sounds were recorded along a transect at 10, 30, 65, and 100 m from the speaker with a Audio-Technica ATR6550X microphone and a Zoom H4n Pro recorder (44.1 kHz sampling rate, 16 bit amplitude resolution, WAV format). Both the microphone and speaker were placed horizontally at 2 m above ground, attached to vertical poles. Playbacks were conducted at three locations within the park. At each location, we broadcast sounds over one transect in habitat with open understory (a cleared walking path with no vegetation between the speaker and microphone) and an adjacent transect in habitat with closed understory (regular vegetation between the speaker and microphone). The order of the transects (open and closed) was alternated.

#### Synthesized sounds

We used the baRulho function *synth_sounds* to create synthesized sounds that varied in four structural features: frequency (20 values between 0.5 and 10 kHz, every 0.5 kHz), duration (0.1 s and 0.2 s), frequency modulation (pure tones versus frequency modulated sounds, simulated with a brownian bridge motion stochastic process), amplitude modulation (flat amplitude envelopes versus two amplitude peaks with a value 4 times that of the lowest amplitude). We synthesized sounds representing all possible combinations of signal structure with the four varying features, which resulted in 160 unique sounds. Each structure was replicated three times for a total of 480 sounds in the master sound file.

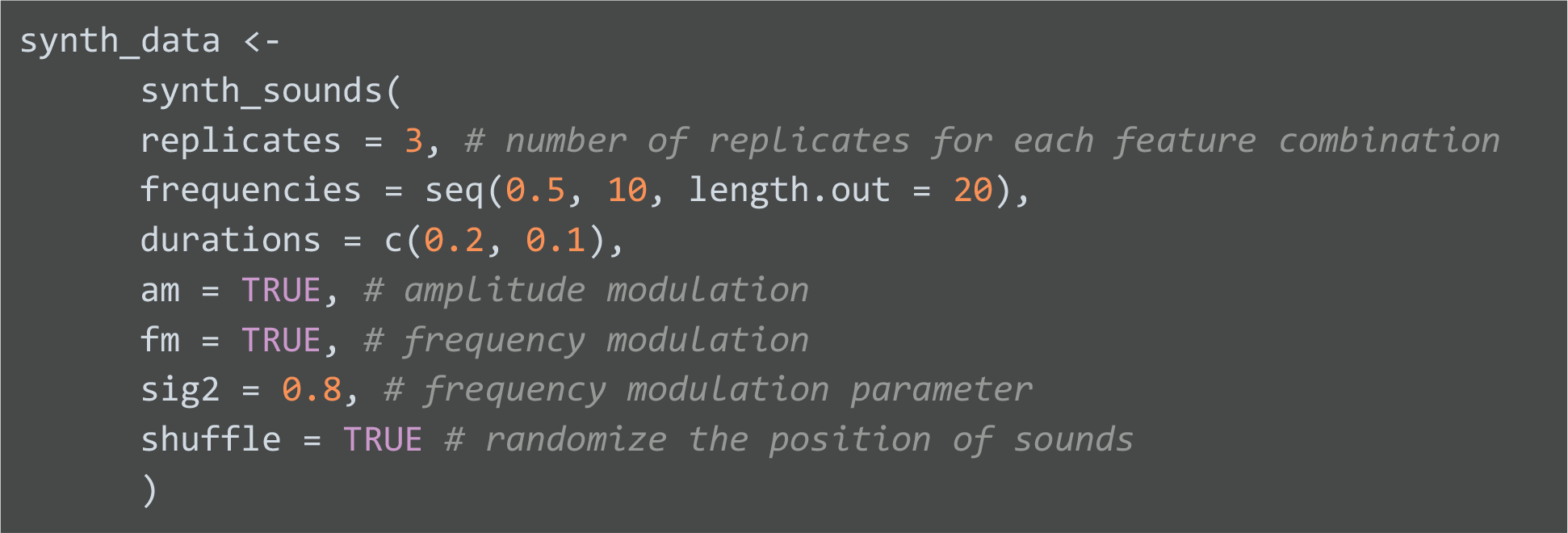

The position of sounds in the master sound file was randomized to avoid adjacent replicates (argument *shuffle =* TRUE). This helps to avoid clustering samples for a given treatment within the master file, which may be affected by fluctuations or irregularities of ambient sounds that mask test sounds during playback experiments. The amount of modulation can be adjusted with the argument *sig2*, which controls the magnitude of frequency change at each step in the brownian motion process. The output of *synth_sounds()* can be used by the function *master_sound_file()* to create a master sound file for playback experiments (i.e. a sound file with all the model sounds in which transmission will be quantified). This function concatenates all sounds into a single sound file, adding silence between sounds (argument *gaps*). In addition, the function adds two acoustic markers at the start and end of the sound files to allow for automated annotation of the re-recorded test files (hereafter test files, explained below; FIG W):

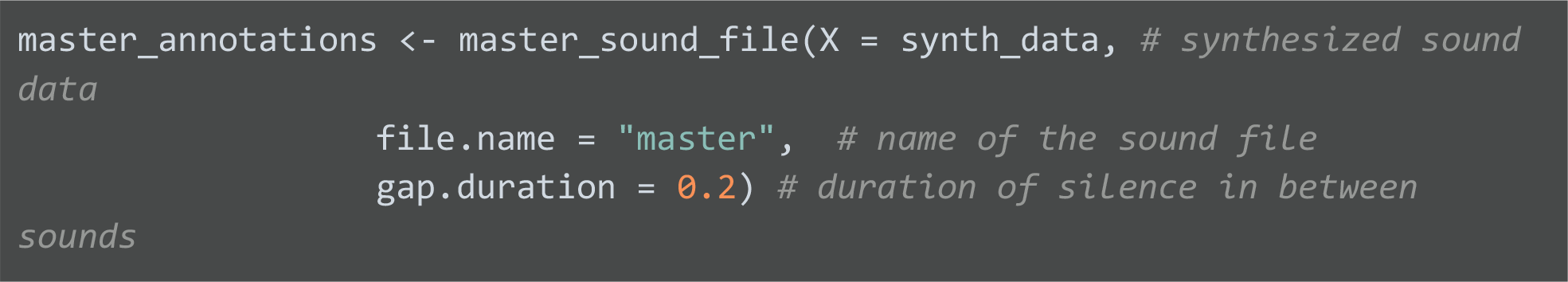

The result is a WAV file added to the current working directory and a data frame with annotations (i.e. time-frequency location) of sounds in the WAV file. The *master_sound_file* function normalizes the amplitude to ensure all sounds have the same maximum amplitude. This function can also be used to create master sound files from annotated animal (non-synthesized) sounds.

#### Time-sync test files

Following playback experiments, the next step is to accurately determine the time location of each test sound among recordings. Without proper alignment, time mismatches between test and reference sounds can produce inaccurate results. Traditionally, the position of test sounds has been determined by visual inspection of waveforms, which can be prone to user error. The package baRulho offers several functions to align recordings in a more systematic way. The function *find_markers* uses spectrographic cross-correlation to locate, in test files, the start and end acoustic markers added by *master_sound_file*. Because the time difference between sounds remains constant, the position of sounds in the test files can be inferred by determining the position of any of the sounds. In this case, the structure of the start and end markers, with higher amplitude (twice that of test sounds by default) and low frequency (2 kHz median frequency by default) make them less prone to degradation during transmission and therefore more likely to be accurately located after being re-recorded. The following code finds acoustic markers over all test sounds from our experiment, which have been shared in the supplementary materials:

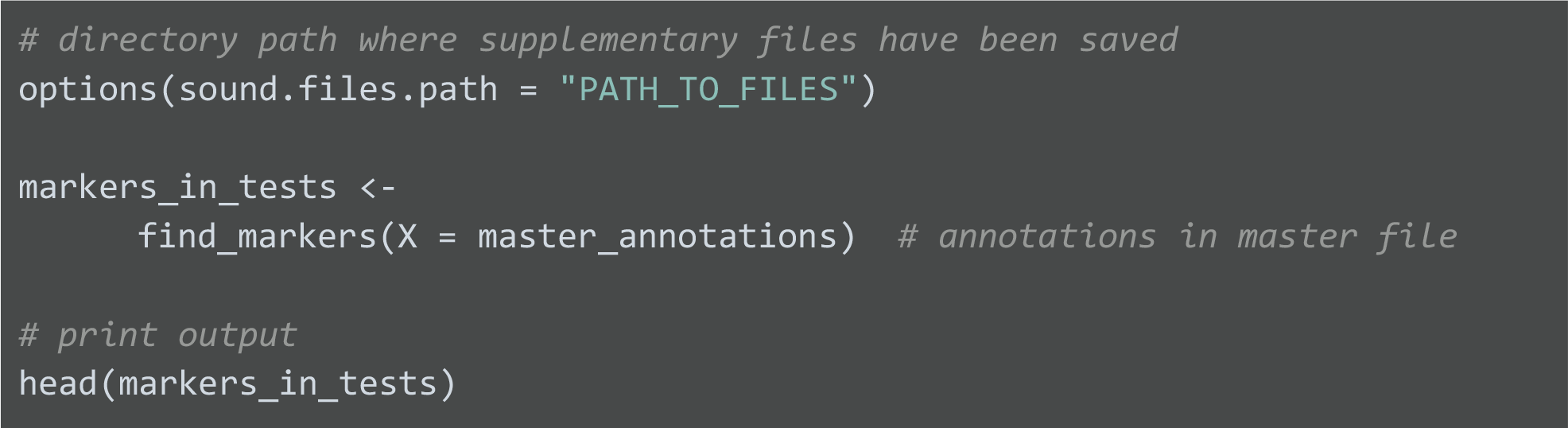

print output head(markers_in_tests) ```
| sound.files | selec | start | end | scores | marker | time.mismatch |
| --- | --- | --- | --- | --- | --- | --- |
| trnsc1_100m_closed.wav | 1 | 6.049203 | 7.049203 | 0.2227792 | start_marker | 0.0300544 |
| trnsc1_100m_closed.wav | 2 | 257.700686 | 258.700686 | 0.2726756 | end_marker | NA |
| trnsc1_100m_open.wav | 3 | 29.502300 | 30.502300 | 0.7150524 | start_marker | 0.0131017 |
| trnsc1_100m_open.wav | 4 | 281.136830 | 282.136830 | 0.6083450 | end_marker | NA |
| trnsc1_10m_closed.wav | 5 | 39.452253 | 40.452253 | 0.7046298 | start_marker | 0.0114997 |
| trnsc1_10m_closed.wav | 6 | 291.085182 | 292.085182 | 0.8364759 | end_marker | NA |

The output contains time-frequency coordinates of the start and end markers for each test file. In addition, the output also contains a ‘time.mismatch’ column that compares the time gap between the two markers in the test files against that in the master sound file. For perfect detection the value must equal 0, therefore, this number can be used as a measure of the maximum possible error. The output from *find_markers* can then be used by the function *align_test_files* to determine the location of all other test sounds:

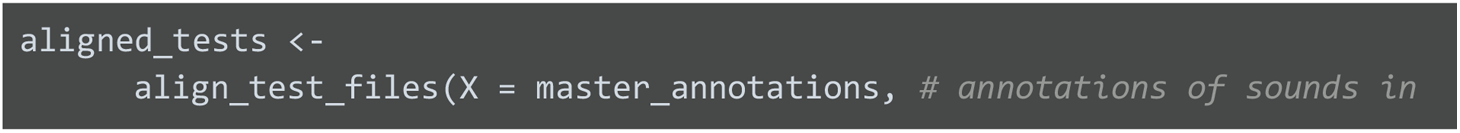

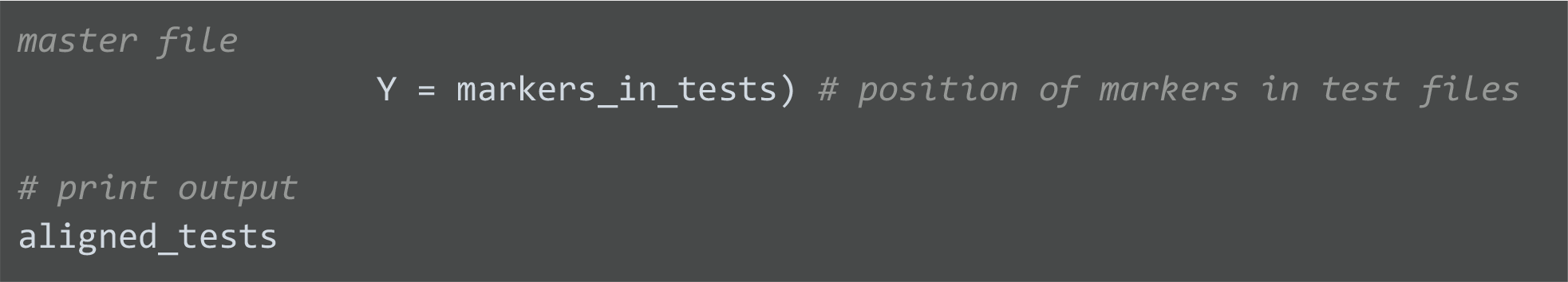

| sound.files | selec | start | end | bottom.freq | top.freq | sound.id | marker |
| --- | --- | --- | --- | --- | --- | --- | --- |
| trnsc1_100m_closed.wav | 1 | 7.129257 | 7.329280 | 7.875 | 8.805 | dur:0.2;freq:9;fm;am_1 | end_marker |
| trnsc1_100m_closed.wav | 2 | 7.379280 | 7.579303 | 3.208 | 4.069 | dur:0.2;freq:4;fm;am_1 | end_marker |
| trnsc1_100m_closed.wav | 3 | 7.629303 | 7.829325 | 0.422 | 1.223 | dur:0.2;freq:1;fm;am_1 | end_marker |
| trnsc1_100m_closed.wav | 4 | 7.879325 | 8.079348 | 6.905 | 7.917 | dur:0.2;freq:7;fm;am_1 | end_marker |
| trnsc1_100m_closed.wav | 5 | 8.129348 | 8.329371 | 8.417 | 9.416 | dur:0.2;freq:9.5;fm;am_1 | end_marker |
| trnsc1_100m_closed.wav | 6 | 8.379371 | 8.579393 | 4.171 | 4.839 | dur:0.2;freq:4.5;fm;am_1 | end_marker |

The function *align_test_files* uses the marker with the highest cross-correlation to estimate the location for all other sounds in the test files. In cases where the location of test sounds (i.e. individual sounds in test files) is still slightly off, sounds can be further aligned with the functions *auto_realign* and *manual_realign* (Table 1). The output of *align_test_files*, (and from *auto_realign* and *manual_realign*) contains the time-frequency coordinates for all test sounds (Fig. 1). The precision of the alignment can be visually inspected using the function *plot_aligned_sounds*, which produces images similar to the spectrograms in each individual panel in Figure 1.

**Figure 1.**
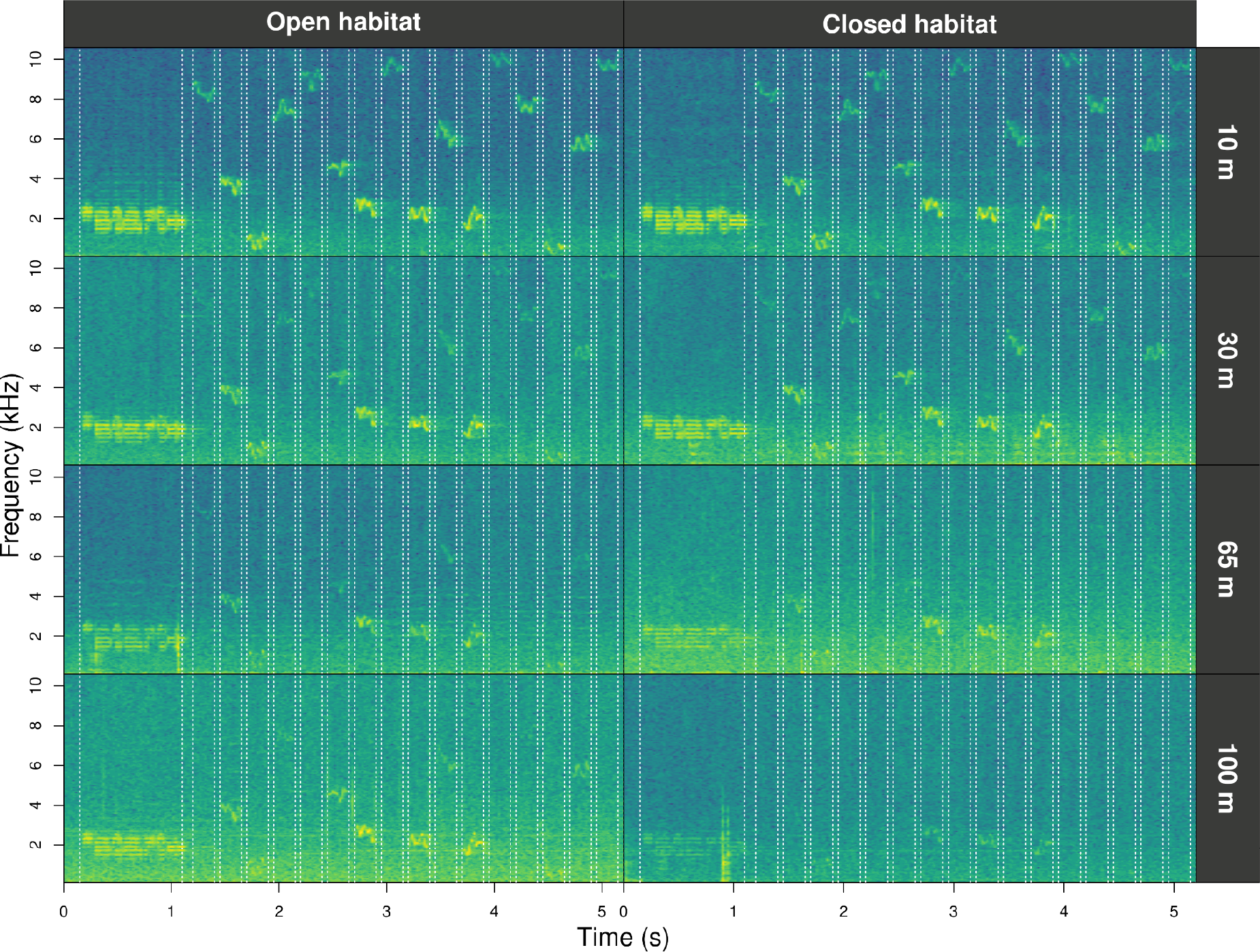
Fourier spectrograms of test recordings from an experimental transect in open and closed habitats (columns) re-recorded at four distances from the playback speaker (rows). The dotted vertical lines highlight the detected position of sounds computed by the functions find_markers and align_test_files.

Alignments can be further adjusted manually with the function *manual_realign*. The function generates a multipanel graph with a spectrogram of the master sound file on top of that from test sound files. This highlights the position of correspondent test sounds on both in order to assess and adjust alignment. The lower spectrogram shows a series of ‘buttons’ that users can click on to control the time position of test sound files:

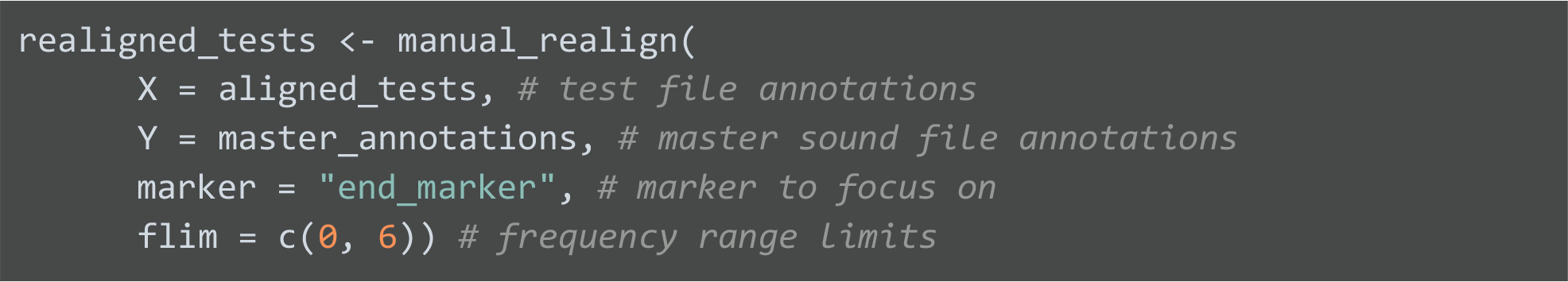

The function returns an object similar to the input test file annotations in which the start and end of the sounds have been adjusted based on user input. The format of annotations is shared by other bioacoustics R packages, which enables the use of additional functionalities, like exporting annotations to external software for further inspection (Rraven, Araya-Salas 2017) or signal structure quantification and additional visualizations (warbleR; Araya-Salas & Smith-Vidaurre 2017).

The last stage of analysis involves the quantification of sound degradation. Most degradation measures require a comparison between test sounds that were recorded at different distances from the speaker, to their reference, which is often re-recorded at 1m. Hence, a column that indicates the distance at which each sound was recorded is needed. The function *set_reference_sound*s determines annotation to be used as reference for each test sound:

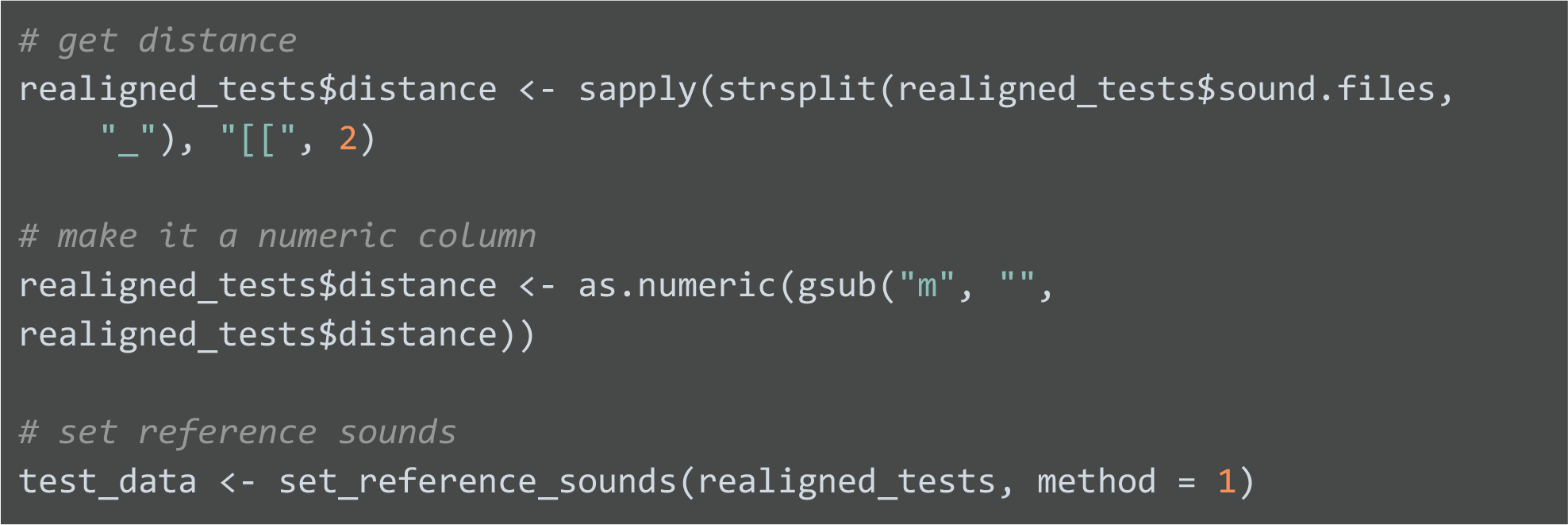

If the default method is used (‘method = 1’), the reference is determined as the re-recorded sample of a sound recorded at the shortest distance. The function adds a column “reference” that is used by functions for measuring degradation. We use the output of this function to compute all eight degradation measures available in baRulho (Table 1). Here, we present the code for four such measures to show how easily they can be obtained with baRulho:

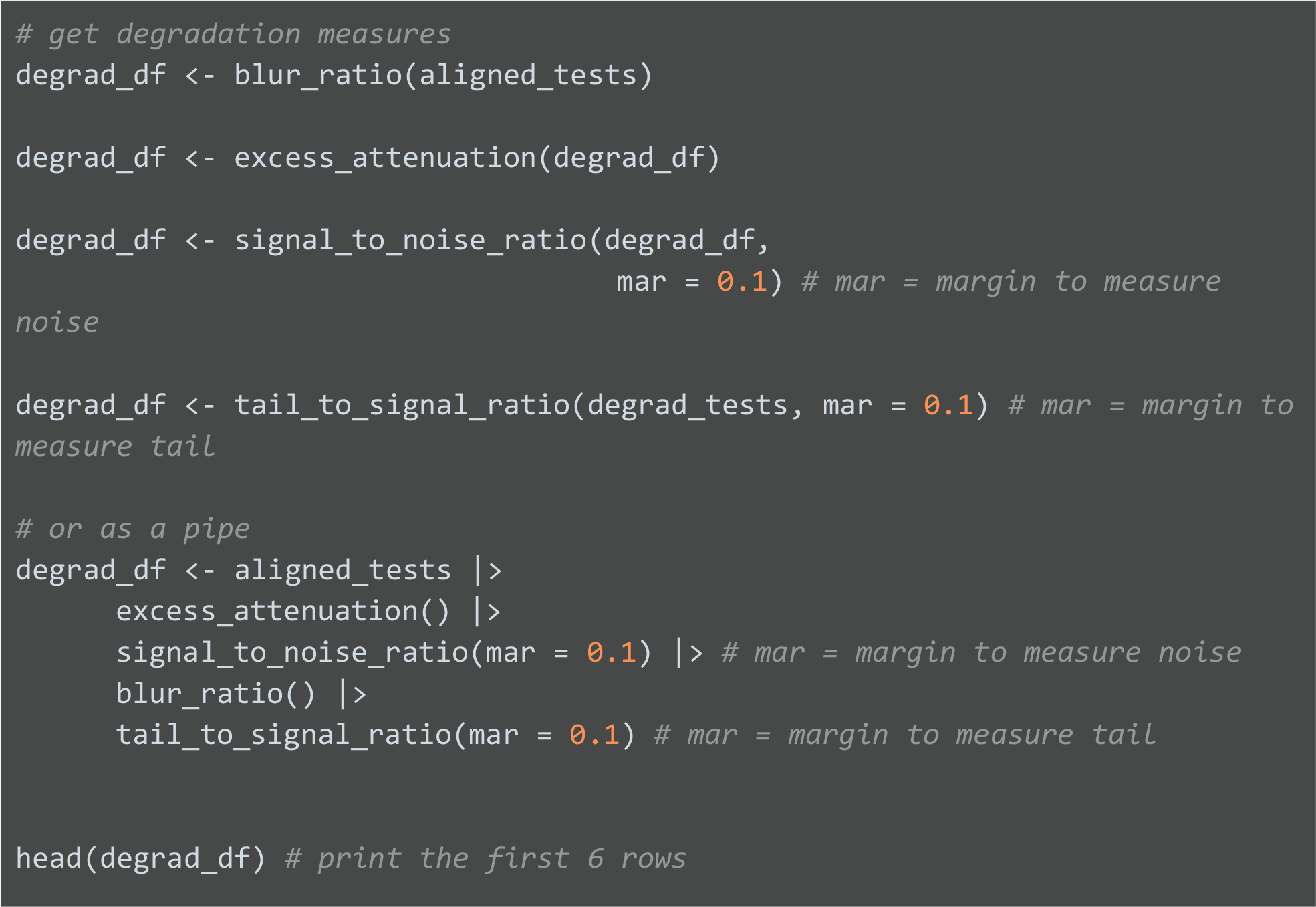

| sound.files | selec | sound.id | distance | reference | excess.attenua<br>tion | signal.to.nois<br>e.ratio | blur.ratio | tail.to.signal.r<br>atio |
| --- | --- | --- | --- | --- | --- | --- | --- | --- |
| trnsc1_100m_closed.wav | 1 | dur:0.2;freq:9;fm;am_1 | 100 | trnsc1_1m_o<br>pen.wav-2 | -10.964.408 | 0.7691817 | 0.2911726 | -0.9995171 |
| trnsc1_100m_closed.wav | 2 | dur:0.2;freq:4;fm;am_1 | 100 | trnsc1_1m_o<br>pen.wav-3 | -1.079.939 | 0.6385018 | 0.2971347 | 0.1901690 |
| trnsc1_100m_closed.wav | 3 | dur:0.2;freq:1;fm;am_1 | 100 | trnsc1_1m_o<br>pen.wav-4 | -7.309.884 | 45.278.465 | 0.1437131 | 24.441.865 |
| trnsc1_100m_closed.wav | 4 | dur:0.2;freq:7;fm;am_1 | 100 | trnsc1_1m_o<br>pen.wav-5 | -11.257.276 | -13.819.294 | 0.2678079 | 0.0593048 |
| trnsc1_100m_closed.wav | 5 | dur:0.2;freq:9.5;fm;am_1 | 100 | trnsc1_1m_o<br>pen.wav-6 | -2.927.407 | -0.2573796 | 0.2060300 | -12.052.163 |
| trnsc1_100m_closed.wav | 6 | dur:0.2;freq:4.5;fm;am_1 | 100 | trnsc1_1m_o<br>pen.wav-7 | -2.767.117 | -0.7123177 | 0.2216715 | 16.697.592 |

The output includes measures of degradation for each test sound (time and frequency position columns are excluded for simplicity). Measurements that involve a comparison between test and reference sounds do not return a value for the reference sounds (recorded at 1m). All functions compute degradation measures within the frequency range of model sounds provided in the annotations used for creating the master sound file. To analyze transmission patterns, we constructed a measure of “overall degradation” based on all eight degradation measures (Table 1), computed as the first component of a Principal Component Analysis on zero-mean, unit variance degradation measures.

Sound degradation at increasing distances can be visually inspected with the function *plot_degradation*. This function produces JPEG files with a mosaic of visual representations of sounds (Fourier spectrograms, power spectrum, and amplitude envelopes) for each test sound and corresponding reference sound (Fig. 2):

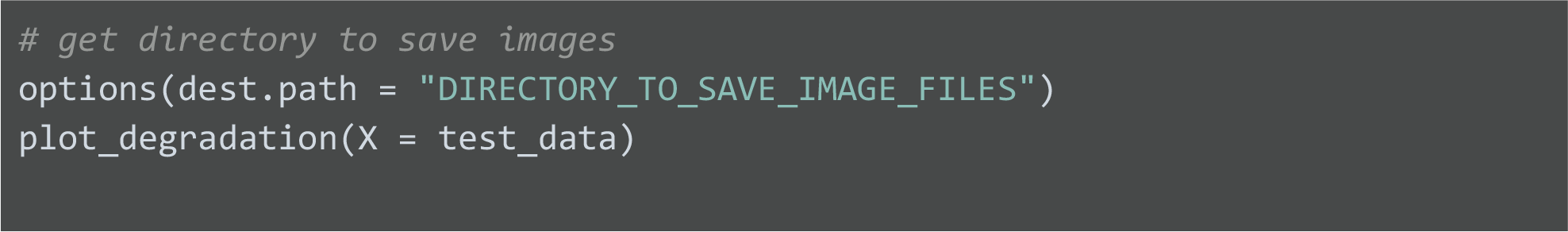

**Figure 2.**
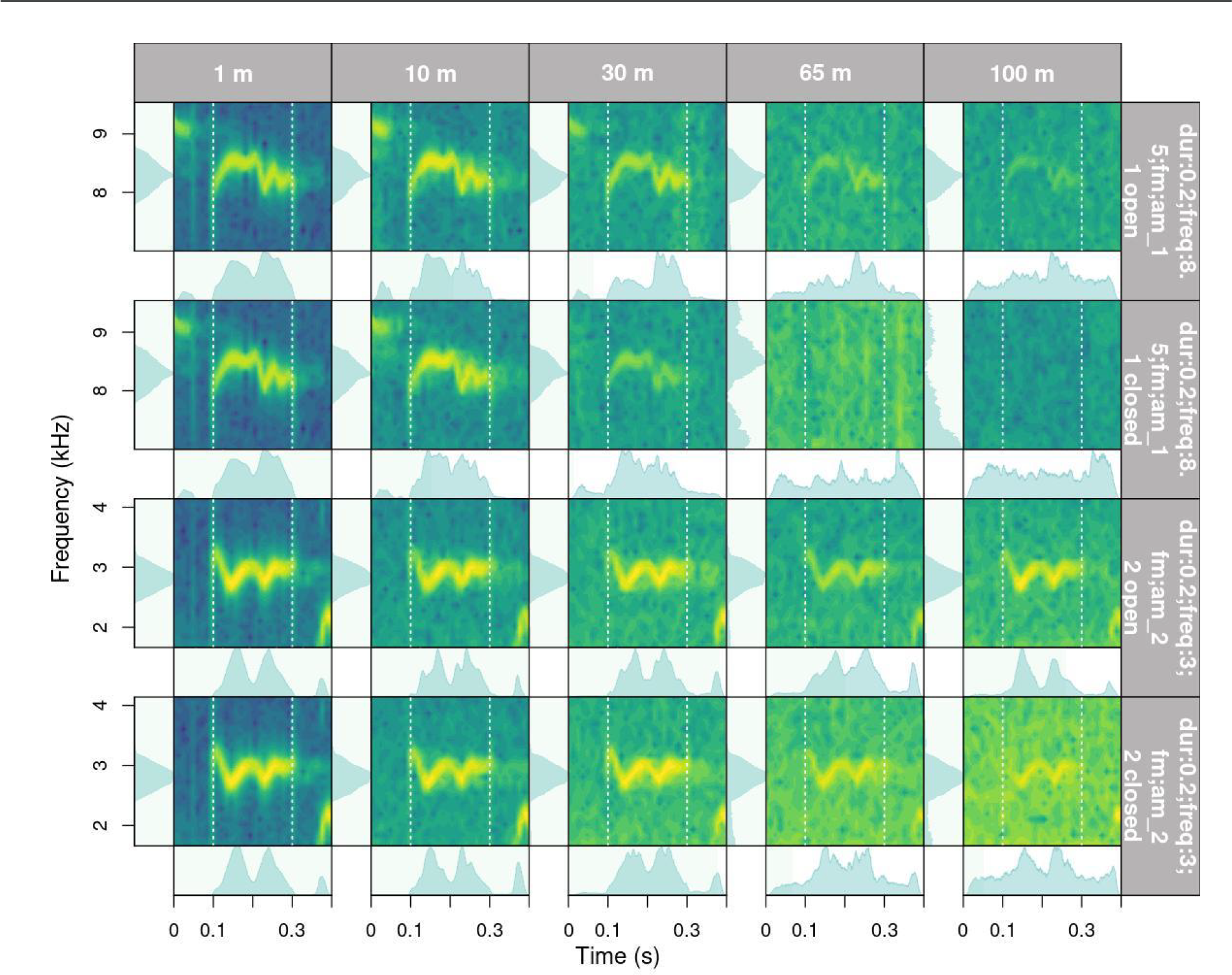
Output from function plot_degradation that shows the Fourier spectrogram, amplitude envelope, and power spectrum for test sounds (distances 10, 30, 65 and 100 m) and their corresponding reference sound (1 m). The first and second row contain a 0.2 s long, 8.5 kHz, frequency and amplitude modulated sound recorded in open and closed understory, respectively. The third and fourth row contain a 0.2 s long, 3 kHz, frequency and amplitude modulated sound recorded in open and closed understory.

The function *plot_blur_ratio* can also be used to visually inspect degradation, which creates JPEG image files with spectrograms of the reference and test sounds (one image for each test sound) and the overlaid power distribution (either amplitude envelopes or power spectrum, see argument ‘type’) as probability mass functions (Fig. 3). The output graphs highlight the mismatch between the two compared distributions (reference and test), which represents the computed blur ratio returned by either *blur_ratio* or *spectrum_blur_ratio*.

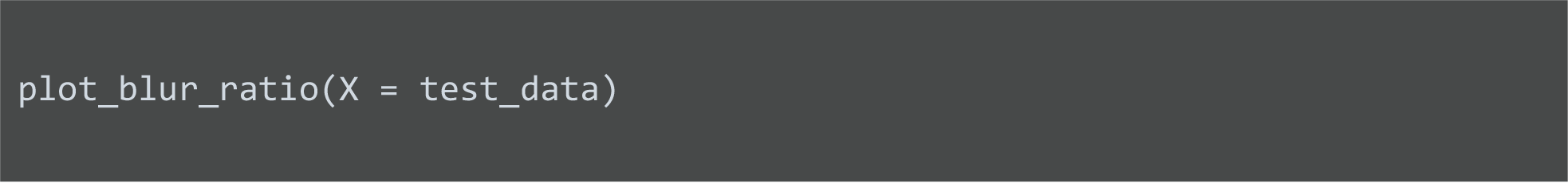

**Figure 3.**
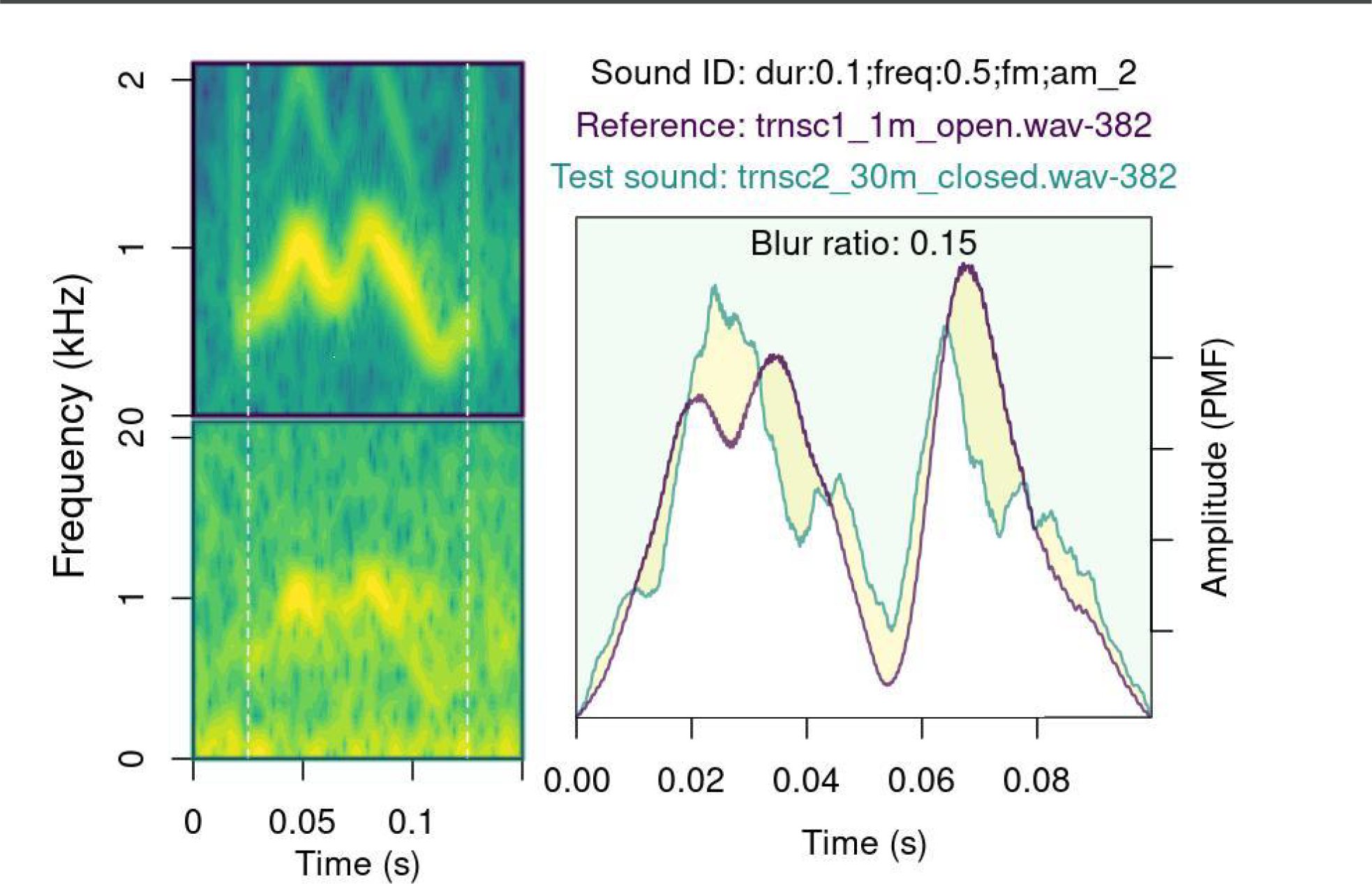
Output from plot_blur_ratio that shows spectrograms of the reference and test sound, the overlaid amplitude envelopes, and the mismatch between the two (yellowish area between the reference [purple line] and test sound [green line]).

### 2.2 Statistical Analysis

#### Patterns of signal degradation

We used Bayesian regression models to evaluate the effect of sound structural features (frequency, frequency modulation, amplitude modulation, and duration) and habitat structure on four degradation measures: blur ratio, excess attenuation, tail-to-signal ratio, and overall degradation (PC1). Models included interaction terms between habitat structure and each of the sound structural features. Distance was modeled as a monotonic effect in which distance levels are not assumed to be equidistant with respect to their effect on the response variable, rather are estimated from the data (Bürkner & Charpentier 2020). Frequency was mean-centered and scaled by dividing it by two standard deviations to allow direct comparisons between effect sizes from categorical predictors (Schielzeth 2010; Gelman 2008). Single response regressions were run for each of the degradation measures. Regressions were run in Stan (Stan Development Team 2021) through the R package brms (Bürkner 2017). Response variables were modeled with a normal distribution. Transect and sound replicates were included as varying intercept effects to account for the paired nature of transects and the non-independence of observations, respectively. Effect sizes are presented as median posterior estimates and 95% credibility intervals as the highest posterior density interval. Minimally informative priors were used for population-level effects (normal distribution; mean = 0; SD = 2) and their standard deviation (Cauchy distribution; median = 0; scale = 4). Predictors in which credible intervals did not include zero were regarded as having an effect on the response variables. Models were run on four chains for 5000 iterations, following a warm-up of 5000 iterations. The effective sample size was kept above 1500 for all parameters. Performance was checked visually by plotting the trace and distribution of posterior estimates for all chains. We also plotted the autocorrelation of successive sampled values to evaluate the independence of posterior samples, ensured that the potential scale reduction factor for model convergence was kept below 1.01 for all parameter estimates and generated plots from posterior predictive samples to assess the adequacy of the models in describing the observed data (Supplemental Materials).

### 2.3 Comparing baRulho and Sigpro

Sigpro (Pedersen 1998) is, to our knowledge, the only software package specifically dedicated to quantifying animal sound degradation (Holland et al. 1998; Balsby et al. 2003; Lampe et al. 2007; Darden et al. 2008; Barker et al. 2009; Graham et al. 2017; Vargas-Castro et al. 2017; Wheeldon et al. 2022). Therefore, we compared measurements extracted in baRulho and Sigpro to evaluate their equivalence. In both programs, we noted the amount of time to complete each analysis. Measurements were taken on a subset of the recording data that included 80 test sounds (20 sounds re-recorded at 4 distances). For our comparison, we used four degradation measures that are included in both software packages: blur-ratio, excess attenuation, signal-to-noise ratio, and tail-to-signal ratio. The analysis was run by MQO, who was already trained on both packages, on a laptop computer (Lenovo, 16 GB RAM memory, Intel Core i7-1255U (1.70 GHz)). In Sigpro, we tested sounds in two ways, first with the original test files, which is the default procedure in Sigpro. This required manually setting the time location for test sounds within the entire re-recorded sound file. In some cases manual placement is difficult, especially at the furthest recording distances where some signals leave almost no trace on the spectrograms, which potentially increases measurement error. Hence, we ran an additional analysis in Sigpro on individual audio clips for each test sound, in which the start of the clip matched the time location used in baRulho. We expected this approach would minimize measurement error, providing a more direct comparison between the two software packages. We used Pearson correlation to quantify agreement between degradation measures in Sigpro and baRulho.

## 3. Results & Discussion

The baRulho package streamlines the process of evaluating acoustic signal degradation, including tasks that previously lacked programmatic solutions. Particularly relevant is the functionality related to simulating model sounds with animal-like attributes for transmission playbacks, concatenating annotated sounds into master sound files for playback experiments, and automating the location of test sounds in the re-recorded sound files. The package also introduces two new degradation measures (i.e. spectrum_correlation and spectrum_blur_ratio) to explore and quantify patterns of sound propagation. Automated annotation, parallel computing, and seamless integration of data batch processing throughout the entire workflow enhances computational efficiency, facilitating analysis of large datasets.

Measures incorporated in the baRulho package adequately quantify acoustic degradation, as the results of our case study are consistent with the observed patterns of sound degradation in natural environments. We found that habitat structure (i.e. open or closed understory) was a primary driver of sound degradation (Fig. 4): degradation in closed understory was more pronounced than in open understory. This effect was evident on the first principal component (overall degradation) as well as on three models that tested single degradation measures (blur ratio, excess attenuation, and tail-to-signal ratio; Fig. 4). In closed habitats, vegetation impedes signal propagation, often through scattering, which results in higher distortion (blur ratio) and reverberation (tail-to-signal ratio) (Dabelsteen et al. 1993; Balsby et al. 2003). Leafy vegetation may also absorb signals as they propagate through closed habitat, and as a result, leads to additional loss of signal intensity or excess attenuation (Holland et al. 1998; Darden et al. 2008).

**Figure 4.**
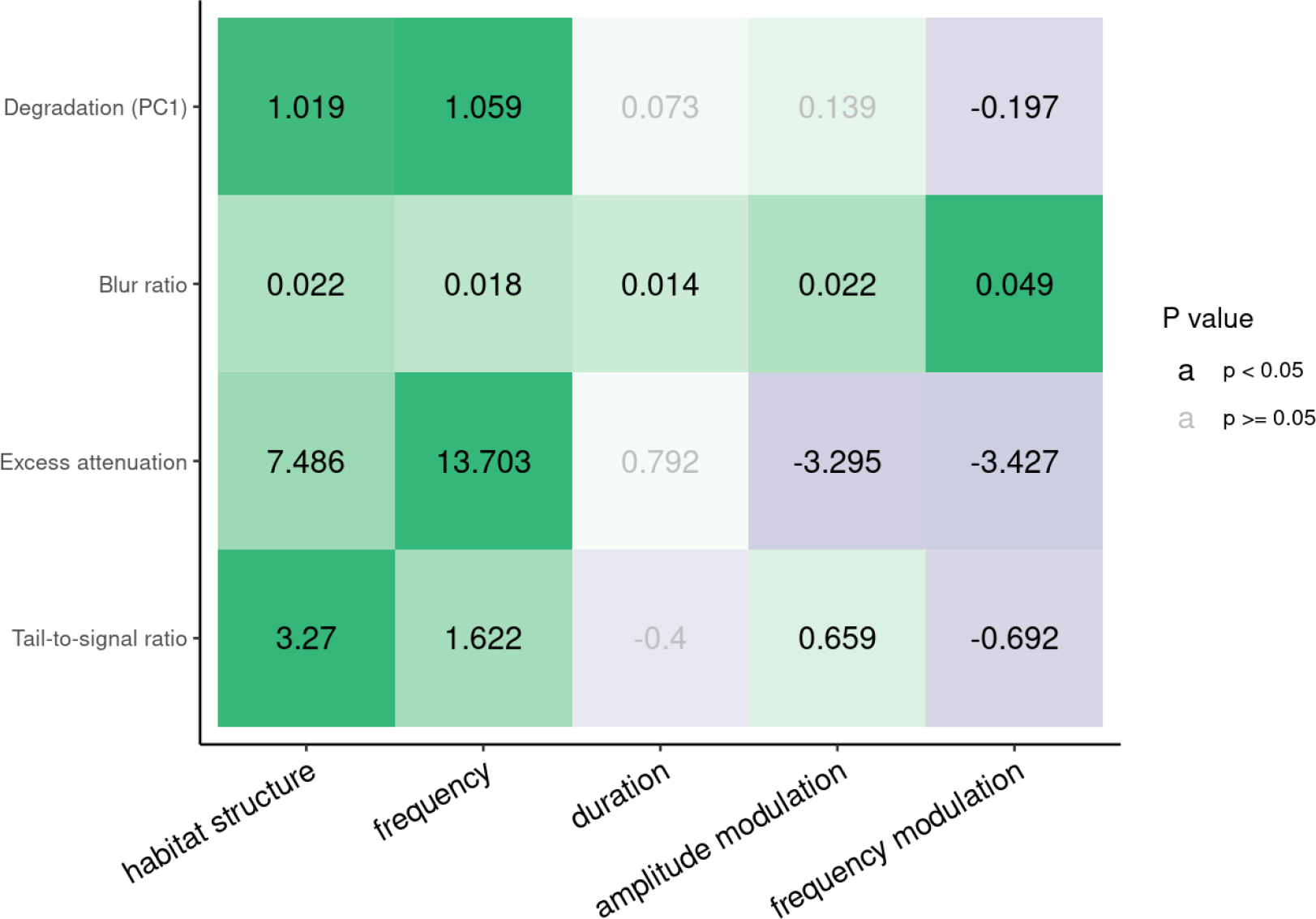
Effect size estimates of predictor variables representing habitat and signal structure (columns) on the signal degradation response variables (rows). Values in gray indicate effects that did not differ from zero. Single models were run for each response. The cell color gradient denotes the magnitude of the effects relative to the highest effect size within a model. Green and purple colors indicate positive and negative effects, respectively.

Frequency was the structural feature more closely linked to degradation (Fig. 4). Higher frequencies show greater overall degradation, excess attenuation, reverberation, and, to a minor degree, amplitude envelope blurring (Fig. 4). Low frequency sounds, due to longer wavelengths, are less susceptible to scattering, thus attenuate less over distance compared to higher frequency sounds (Wiley & Richards 1982). Frequency modulation and, to a lesser extent amplitude modulation, contributed to distortion of the amplitude envelope (blur ratio; Fig. 4). Tonal sounds withstand scattering better than frequency modulated sounds (Morton 1975; Marten & Marler 1977). Accordingly, low frequency, tonal sounds are often considered advantageous for long-distance communication in closed habitats, such as forests, as salient information may be better detected over larger distances (Yip et al. 2017). However, we also found differences in the way each structural feature influenced degradation. Particularly relevant is the lower degradation, attenuation and reverberation of frequency modulated sounds compared to pure tones. Amplitude modulated sounds were also less attenuated than their non-modulated counterparts (Fig. 4). Over distance, original patterns of amplitude modulation elicited from a sender declines or blurs (Dabelsteen et al. 1993; Apol et al. 2017). Nonetheless, amplitude modulated sounds are more detectable at lower signal-to-noise ratios than sounds without amplitude modulation (Lohr et al. 2003), which could explain the observed pattern. Overall, our results are consistent with predictions from the Acoustic Adaptation Hypothesis (AAH), which posits that habitat-dependent degradation imposes a strong selective pressure on animal acoustic signals, favoring features that optimize efficient transmission (Morton 1975).

Overall, we found agreement between baRulho and Sigpro results. Measurement agreement was modest between the two software when we manually located test sounds in Sigpro (tail-to-signal ratio: r = 0.61; signal-to-noise ratio: r = 0.79; blur ratio: r = 0.5; excess attenuation: r = 0.75). However, agreement improved markedly when Sigpro measures were taken on recording clips that started at the onset of sounds, which we determined in baRulho (tail-to-signal ratio: r = 0.8; signal-to-noise ratio: r = 0.94; blur ratio: r = 0.8; excess attenuation: r = 0.97; Fig. 5). The observed agreement indicates that degradation measures obtained with baRulho are comparable to those from Sigpro. The observed improvement when time location between recordings for the Sigpro analysis was determined using the baRulho approach suggests that much of the mismatch between the two software packages is due to the way sound position is determined in each program. Sigpro requires visual inspection of the waveform to manually identify test and reference sounds, which must be repeated even when the same reference is used several times. In contrast, baRulho employs spectrogram cross-correlation to find the position of acoustic markers, which are then used to compute positions for all other sounds (Fig. 1). Furthermore, baRulho provides a means for identifying poor location estimates and additional functionality to improve their accuracy. Spectrogram cross-correlation for signal alignment is beneficial, given that this approach successfully determined the position of all test sounds, even on recordings made 100 m from the speaker. Our package also represents a significant improvement in computational efficiency. The analysis of the subset of 80 test sounds in Sigpro took 861 min to complete. In contrast, the same analysis in baRulho took 7.43 minutes (26.65 seconds of computing time running the script using 8 cores), on a regular laptop computer, which is approximately 119 times faster.

**Figure 5.**
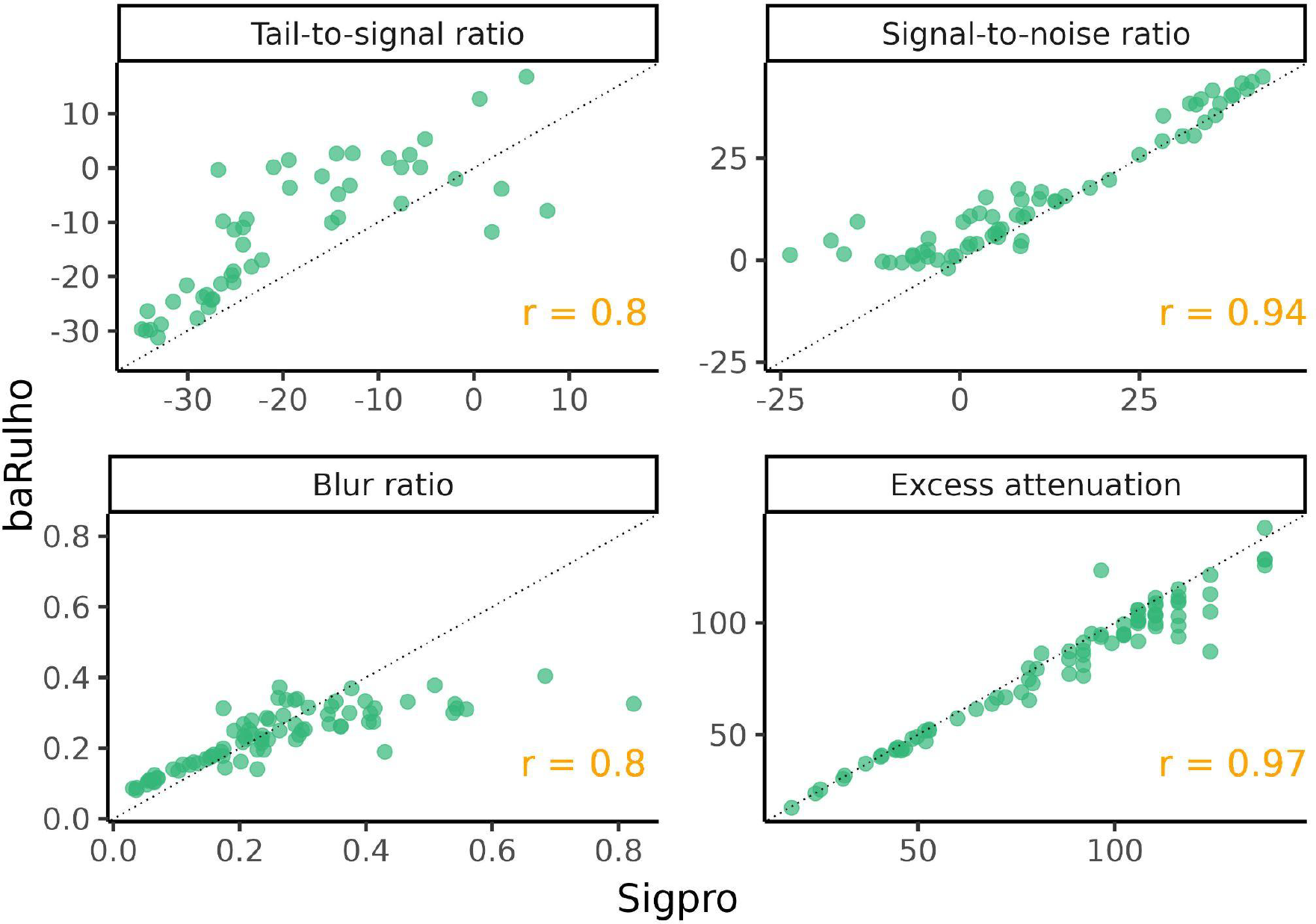
Scatterplots showing the agreement between baRulho and Sigpro for the four degradation measures in common. The Pearson correlation coefficient for each comparison is shown in the bottom right corner of each panel. Sigpro measures shown were taken on recording clips that started at the onset of sounds, which was determined in baRulho.

## 4. Conclusions

The R package baRulho provides a user-friendly suite of tools designed to measure animal sound degradation via signal transmission experiments. We show how the package can be used across the different stages of a transmission experiment, from data preparation to quantification of degradation. The sound analysis results generated by baRulho and Sigpro are similar, making results comparable across studies using either software. However, baRulho provides significant improvements to previous approaches as manual annotation is not required to determine test sound location, much less time is needed to complete data extraction, new measures are added to quantify novel aspects of degradation, and new visualizations are available for inspecting degradation. We expect this package to be a useful contribution to the community that can enable researchers to bring more detailed insight into the evolutionary processes that have shaped animal acoustic communication in natural environments.

## Acknowledgements

We thank Luis Sandoval (Universidad de Costa Rica) for guidance on early stages of package development and participants of the workshop on Bioacoustic Analysis in R at Instituto de Biología, Universidad Nacional Autónoma de México for assistance during field work. A.R-G. is thankful for support through the Walt Halperin Endowed Professorship and the Washington Research Foundation as Distinguished Investigator.

## Conflicts of Interest

The authors declare no conflicts of interest.

## Data Availability Statement

The baRulho package is available on CRAN

(https://cran.r-project.org/package=baRulho). The development version of the package used in the application and the source code can be found at https://github.com/maRce10/baRulho. The scripts and data for running the example code in the paper are available at https://github.com/maRce10/barulho_paper and https://doi.org/10.6084/m9.figshare.21559074.v3, respectively.

## Notes

https://marce10.github.io/baRulho/

https://rpubs.com/marcelo-araya-salas/1075886

https://doi.org/10.6084/m9.figshare.21559074.v3

### Competing Interest Statement

The authors have declared no competing interest.

## References

Apol, C. A., Sturdy, C. B., & Proppe, D. S. (2018). Seasonal variability in habitat structure may have shaped acoustic signals and repertoires in the black-capped and boreal chickadees. Evolutionary Ecology, 32, 57–74.

Arasco, A. G., Manser, M., Watson, S. K., Kyabulima, S., Radford, A. N., Cant, M. A., & Garcia, M. (2022). Testing the acoustic adaptation hypothesis with vocalizations from three mongoose species. Animal Behaviour, 187, 71–95

Araya-Salas, M. (2017). Rraven: connecting R and Raven bioacoustic software. R package version, 1(2).

Araya-Salas, M., & Smith-Vidaurre, G. (2017). warbleR: an R package to streamline analysis of animal acoustic signals. Methods in Ecology and Evolution, 8(2), 184–191.

Balsby, T., Dabelsteen, T., & Pedersen, S. B. (2003). Degradation of whitethroat vocalisations: implications for song flight and communication network activities. Behaviour, 140(6), 695–719

Barker, N. K., Dabelsteen, T., & Mennill, D. J. (2009). Degradation of male and female rufous-and-white wren songs in a tropical forest: effects of sex, perch height, and habitat. Behaviour, 1093-1122.

Benedict, L., Hardt, B., & Dargis, L. (2021). Form and Function Predict Acoustic Transmission Properties of the Songs of Male and Female Canyon Wrens. Frontiers in Ecology and Evolution, 9, 797. 10.3389/fevo.2021.722967

Bradbury, J. W., & Vehrencamp, S. L. (2011). Principles of animal communication, 2nd edn Sunderland. MA: Sinauer Associates.

Brown, T. J., & Handford, P. (2003). Why birds sing at dawn: the role of consistent song transmission. Ibis, 145(1), 120–129. 10.1046/J.1474-919X.2003.00130.X

Bürkner, P.-C. (2017). brms: An R Package for Bayesian Multilevel Models Using Stan. Journal of Statistical Software, 80(1), 1–28. doi:10.18637/jss.v080.i01

Bürkner, P.-C. and Charpentier, E. (2020). Modelling monotonic effects of ordinal predictors in Bayesian regression models. Br J Math Stat Psychol, 73: 420–451. 10.1111/bmsp.12195

Cardoso, G. C., & Price, T. D. (2010). Community convergence in bird song. Evolutionary Ecology, 24, 447–461.

Dabelsteen, T., Larsen, O. N., & Pedersen, S. B. (1993). Habitat-induced degradation of sound signals: Quantifying the effects of communication sounds and bird location on blur ratio, excess attenuation, and signal-to-noise ratio in blackbird song. The Journal of the Acoustical Society of America, 93(4), 2206–2220.

Darden, S. K., Pedersen, S. B., Larsen, O. N., & Dabelsteen, T. (2008). Sound transmission at ground level in a short-grass prairie habitat and its implications for long-range communication in the swift fox Vulpes velox. The Journal of the Acoustical Society of America, 124(2), 758–766.

Endler, J. A. (1992). Signals, signal conditions, and the direction of evolution. The American Naturalist, 139, S125–S153

Gelman, A. (2008). Scaling regression inputs by dividing by two standard deviations. Statistics in medicine, 27(15), 2865–2873.

Grabarczyk, E. E., & Gill, S. A. (2020). Anthropogenic noise masking diminishes house wren (Troglodytes aedon) song transmission in urban natural areas. Bioacoustics, 29(5), 518–532.

Graham, B. A., Sandoval, L., Dabelsteen, T., & Mennill, D. J. (2017). A test of the Acoustic Adaptation Hypothesis in three types of tropical forest: degradation of male and female Rufous-and-white Wren songs. Bioacoustics, 26(1), 37–61.

Hardt, B., & Benedict, L. (2020). Can you hear me now? A review of signal transmission and experimental evidence for the acoustic adaptation hypothesis. Bioacoustics, 30(6), 716–742. 10.1080/09524622.2020.1858448

Holland, J., Dabelsteen, T., Pedersen, S. B., & Larsen, O. N. (1998). Degradation of wren Troglodytes troglodytes song: implications for information transfer and ranging. The Journal of the Acoustical Society of America, 103(4), 2154–2166

Kime, N. M., Turner, W. R., & Ryan, M. J. (2000). The transmission of advertisement calls in Central American frogs. Behavioral Ecology, 11(1), 71–83

Lampe, H., Larsen, O., Pedersen, S., & Dabelsteen, T. (2007). Song degradation in the hole-nesting pied flycatcher Ficedula hypoleuca: implications for polyterritorial behaviour in contrasting habitat-types. Behaviour, 144(10), 1161–1178.

LaZerte, S. E., Otter, K. A., & Slabbekoorn, H. (2015). Relative effects of ambient noise and habitat openness on signal transfer for chickadee vocalizations in rural and urban green-spaces. Bioacoustics, 24(3), 233–252.

Leader, N., Wright, J., & Yom-Yov, Y. (2005). Acoustic properties of two urban song dialects in the orange-tufted sunbird (Nectarinia osea). The Auk, 122(1), 231–245.

Lohr, B., Wright, T. F., & Dooling, R. J. (2003). Detection and discrimination of natural calls in masking noise by birds: estimating the active space of a signal. Animal Behaviour, 65(4), 763–777.

Marten, K., & Marler, P. (1977). Sound transmission and its significance for animal vocalization - I. Temperate habitats. Behavioral Ecology and Sociobiology, 2(3), 271–290. 10.1007/BF00299740

Morton, E. S. (1975). Ecological sources of selection on avian sounds. The American Naturalist, 109(965), 17–34.

Naguib, M., Schmidt, R., Sprau, P., Roth, T., Flörcke, C., & Amrhein, V. (2008). The ecology of vocal signaling: male spacing and communication distance of different song traits in nightingales. Behavioral Ecology, 19(5), 1034–1040.

Nemeth, E., Dabelsteen, T., Pedersen, S. B., & Winkler, H. (2006). Rainforests as concert halls for birds: Are reverberations improving sound transmission of long song elements? The Journal of the Acoustical Society of America, 119(1), 620. 10.1121/1.2139072

Pedersen, S. B. (1998). Preliminary operational manual for signal processor Sigpro. Centre of Sound Communication, Odense University, Odense.

Schielzeth, H. (2010). Simple means to improve the interpretability of regression coefficients. Methods in Ecology and Evolution, 1(2), 103–113.

Tobias, J. A., Aben, J., Brumfield, R. T., Derryberry, E. P., Halfwerk, W., Slabbekoorn, H., & Seddon, N. (2010). Song divergence by sensory drive in Amazonian birds. Evolution: International Journal of Organic Evolution, 64(10), 2820–2839. http://dx.doi.org/10.1111/j.1558-5646.2010.01067.x

Vargas-Castro, L. E., Sandoval, L., & Searcy, W. A. (2017). Eavesdropping avoidance and sound propagation: the acoustic structure of soft song. Animal behaviour, 134, 113–121.

Wheeldon A, Kwiatkowska K, Szymański P, Osiejuk TS (2022). Male and female song propagation in a duetting tropical bird species in its preferred and secondary habitat. PLoS ONE, 17(10): \pe0275434. 10.1371/journal.pone.0275434

Wiley, R. H., & Richards, D. G. (1978). Physical constraints on acoustic communication in the atmosphere: implications for the evolution of animal vocalizations. Behavioral ecology and sociobiology, 3, 69–94

Yip, D. A., Bayne, E. M., Sólymos, P., Campbell, J., & Proppe, D. (2017). Sound attenuation in forest and roadside environments: Implications for avian point-count surveys. The Condor, 119(1), 73–84. 10.1650/CONDOR-16-93.1

